# Ecological assessment of the Amazon sailfin catfish (*Pterygoplichthys* species) within the Indian freshwaters: a mesocosm-based approach

**DOI:** 10.1101/2023.07.22.550140

**Authors:** Suman Mallick, Ajmal Hussan, Jitendra Kumar Sundaray, Ratna Ghosal

## Abstract

Amazon sailfin catfish is relatively a recent invader to the open waters of India, and studies on ecological assessment of the species within the Indian freshwaters are lacking. In the present study, to assess the ecological impact of catfish, we established mesocosms mimicking the Indian freshwaters within natural ponds of eastern India using three species of native fish, rohu, catla and mrigal, and for two size classes, small (10-20 cm length) and large (20-30 cm length) native fishes. Mesocosms were maintained with (test) and without the catfish (control), and length and weight of native fish, zooplankton abundance, and several hydrological and soil parameters were measured at a monthly interval for a period of 120-days. The catfish had a significant (P<0.05) negative impact on growth of small-size rohu only. However, we found no significant (P>0.05) differences in abiotic parameters and zooplankton abundance between control and test ponds for the small-size class. We speculate that reduced growth of rohu could be due to competition from catfish in the context of feeding, and not due to modification of abiotic environment. Thus, we emphasize upon the need for behavioral studies to further assess the impact of the catfish.

## INTRODUCTION

Introduction of a species outside its native range can cause irreversible and devastating impact to a natural ecosystem. Such species, commonly referred to as exotics, are mostly introduced via human activities (Simberloff et al., 2013). Globally, freshwater ecosystems are highly threatened by exotic introductions, and this is mostly due to relatively easier transfer of aquatic organisms in comparison to terrestrial species, and lenient regulations on import of aquarium pets. Aquaculture farming has also facilitated introductions of several exotic species in freshwater habitats across the globe. As per FAO, the percentage of commercially farmed exotic species in aquaculture has drastically increased by 31.8% from 2006 to 2018 (Stankus, 2020). In many instances, exotic species go unnoticed for a longer span of time without causing any ecological change in the introduced ecosystems. In contrast, there are instances, where exotic species have caused considerable damage to an introduced ecosystem, including extinction of native species as well as alteration of community structure and ecosystem function, and have ended up being invasive to the given ecosystem. One such example is the extinction of many endemic cichlid fishes in Lake Victoria, Africa, due to deliberate introduction of Nile perch *Lates niloticus* (Cohen, 1994; Pitcher, 1995). The ways in which an exotic will impact a particular ecosystem can only be addressed by systematic assessment of the interactions among the ecological community of native species, the abiotic variables, and the newly introduced, exotic species.

Studies on exotic species have largely focused on detection and abundance rates, and mostly measuring their impact on abiotic factors (Angeler et al., 2003). Several studies have merely relied upon correlating abundance of introduced species with a range of different biotic and abiotic factors, without necessarily conducting any systematic experiments through manipulations (for example, presence or absence of exotics, impact of exotics on different developmental stages of the natives) of different parameters within an ecosystem (Vimos et al., 2015). Manipulative experiments can typically provide greater mechanistic understanding of the drivers of ecological change (Blanchet et al., 2007). Unfortunately, the scope of manipulations under natural conditions are limited and thus, mesocosms provide a great alternative. Mesocosms have the potential to explain biological complexities within the natural systems (Odum, 1984). Odum (1984) regarded mesocosm as bounded and partially enclosed outdoor experimental setups that can bridge the gap between laboratory and wild environment. Further, mesocosms offer a range of manipulations, which are important in the case of exotic species to decipher the exact mechanisms through which an introduced species can possibly alter a given ecosystem (Lähteenmäki et al., 2015; Herrera-Martínez et al., 2017). Apart from systematic understanding, mesocosm experiments are logistically easier to conduct when compared to natural conditions, and thus, can be replicated along spatial and temporal scales, as well.

Each ecosystem is a unique entity (Williamson & Fitter, 1996) and thus, impact assessments of an exotic species need to be conducted across a range of different habitats, particularly for those species that are popular aquarium pets and have had multiple episodes of introductions across the globe. One such popular pet species is our model organism, the Amazon sailfin catfish *Pterygoplichthys* sp. (Wakida-Kusunoki et al., 2007), which has been introduced in several freshwater ecosystems across the globe. Sailfin catfish is native to the streams of Orinoco and Amazon river basins of South America (Page & Robins, 2006). Several anecdotal reports suggest the negative impact of the exotic catfish on the introduced ecosystems. For example, sailfin catfish have been speculated to impact aquatic plant species due to their grazing habits (Hussan et al., 2019), and contribute towards bank erosion in Marikina river, Philippines, as they burrow during the breeding seasons (Jumawan & Herrera, 2014; Hoover et al., 2014). Elfidasari et al., (2020) experimentally showed that post introduction of the Amazon sailfin catfish, at Ciliwung river, Jakarta, Indonesia, the water quality deteriorated in terms of reduced dissolved oxygen (4.66 to 2.6 mg/l), increased alkaline pH (6.5-7.2), increased ammonia concentration (0.6 to 2.65 ppm) and turbidity (18.13-43.85 FTU). Such deterioration of water quality impaired the native community and favoured the sailfin catfish population to flourish.

The sailfin catfish has been introduced in several countries (Levin et al., 2008; Shafland, 1996; Page & Robins, 2006) and have been anecdotally shown to negatively affect the native species, for example, the catfish had displaced several species of minnows in Texas including the threatened Devils River minnow *Dionda diaboli* (Hoover et al., 2014). Studies also speculated that high abundance of the Amazon sailfin catfish contribute towards economic loss as the hard armour of the catfish damages the nets, and diminishes the catch efficiency of the food fish species (Wakida-Kusunoki et al., 2007). Sumanasinghe and Amarasinghe (2013) reported that Amazon sailfin catfish introduced in Pologolla reservoir, Sri Lanka, have possibly reduced catch efficiency of the native cichlids. Though the catfish has been globally introduced in multiple different ecosystems (Ribeiro et al., 2008; Patoka et al., 2020; Hoover et al., 2004) with serious speculations including anecdotal evidences on its detrimental effect on the invaded habitats, a thorough assessment on the impact of the sailfin catfish within the introduced ecosystems is still limited, and thus, needs further investigations.

In this paper, we investigated the impact of the Amazon sailfin catfish using a mesocosm-based approach mimicking the Indian freshwater ecosystem. We measured the impact of catfish over both structural (growth of co-occurring species) and functional components (nutrient dynamics in the form of chlorophyll ‘a’, soil nutrients as well as hydrological parameters) of the mesocosm ponds. Several species of the catfish have invaded peninsular India, and most species hybridize a lot (Jumawan et al., 2011; Chakraborty et al., 2020). Thus, in this paper, we refer to sailfin catfish by including any species that belongs to the *Pterygoplichthys* genus. The occurrence of the Amazon sailfin catfish has been reported from several places that include biodiversity hotspots of Southern India, for example, Vylathur and the Chackai Canal of Kerala (Krishnakumar et al., 2009) and the wetlands of Chennai (Knight, 2010) and have been reported to occur in the Eastern India including Bihar (Sinha et al., 2010) and West Bengal, (Hussan et al., 2016; Suresh et al., 2019) and in the northern part of India, from Andhra Pradesh (Seshagiri et al., 2021) as well as from Uttar Pradesh (Singh, 2014). A few studies have been conducted on impact assessment of catfish in certain ecosystems outside India or under laboratory conditions (Table 1), however, these studies are not exhaustive in nature in terms of including multiple biotic and abiotic factors. Most of these studies either included only one native fish species as the biotic variable or looked at only dietary preferences of native and exotics without any measurements on the abiotic factors. Thus, there seems to be a knowledge gap in mechanistic understanding of the ways in which the introduced sailfin catfish may potentially impact an ecosystem.

**Table 1.** A literature review showing studies that collected empirical data on assessing the impact of sailfin catfish over an ecosystem, including biotic and/or abiotic factors.

| Type of study / Study site | Parameters included | Main results* | Citation |
| --- | --- | --- | --- |
| Mesocosm / Florida, USA | Hydrological parameters (Temperature, conductivity, dissolved oxygen, pH)<br>Growth of native golden topminnow (*Fundulus chrysotus*) | No impact on growth of native topminnow<br>Decline of periphyton due to excessive feeding by the exotic catfish<br>No impact on hydrological parameters in presence of exotic catfish | Hoover et al., 2013 |
| Natural / Mexico, North America | Stable isotope ratios of carbon and nitrogen | Similar isotope signature indicating dietary overlap between native and exotic species. | Mendoza-Carranza et al., 2010 |
| Natural/ Ragama, Sri Lanka | Gut content analysis of exotic catfish and native fish species (8 species of native fishes) | High dietary overlap between exotic catfish and native fish. | Wijethunga & Epa, 2008 |
| Laboratory/ Not Applicable | Weight of native species (only one species) in presence and absence of sailfin catfish. | Exotic catfish had no impact over the growth of native fish. | Chaichana & Jongphadungkiet, 2012 |

\*Objectives and results pertaining to the impact of sailfin catfish on biotic and/or abiotic factors have only been considered.

Based on this background, we used a mesocosm based approach to address the following objectives for our study: a) whether the exotic Amazon sailfin catfish have any impact over the biotic and abiotic factors of the mesocosms mimicking the Indian freshwater ecosystems; b) whether the impact is time dependent, for example, assessing the ecological impact of catfish over time; c) whether the impact varies depending on different size classes of the native fish species. We conducted mesocosm experiments in the inland, natural ponds and selected three species of native fishes, rohu *Labeo rohita*, catla *Labeo catla* and mrigal *Cirrhinus mrigala* for the experiments. All the three native species co-occur in the open water systems along with the introduced catfish (Suresh et al., 2019), and are commonly referred to as Indian Major Carps (IMC). The study provided systematic assessment of the impact of the catfish on multiple biotic and abiotic factors of the mesocosm ponds, and the assessments were conducted for two different size classes (small-size, 10-20 cm, and large-size, 20-30 cm) of the native species.

## MATERIALS AND METHODS

### Study site and mesocosm setup

The study was conducted in inland ponds (0.04 ha) at the field station of the Indian Council of Agricultural Research (ICAR)-Central Institute of Freshwater Aquaculture (CIFA) (22.96°N, 88.44°E), Kalyani, West Bengal, India. The ponds (N=2) were dried and treated with lime (calcium oxide-1000 kg/ha.) before the start of the experiments (Sipaúba-Tavares et al., 2003). Bird net (mesh size of 19 mm) was used to cover the experimental ponds to prevent fish predation by birds. Mesocosm setup for large-size IMC was conducted from November 2020 to March 2021, and that for small-size IMC was from November 2021 to June 2022. For both the size classes, one of the ponds was maintained as control (absence of catfish) and another as test (presence of catfish). Both the control and test ponds (for both the size classes) were stocked with IMC (N=300), namely rohu (*Labeo rohita*), catla (*Labeo catla*), and mrigal (*Cirrhinus mrigala*), following a standard ratio of 1:3:1 (Debnath et al., 2020). We stocked rohu more in the ponds as it has relatively higher economic value when compared to other two carps (Labh et al., 2014). Moreover, in most aquatic ecosystems, rohu acts as a keystone species (Dwivedi et al., 2020). Test ponds had sailfin catfish (N=80), and the size of the catfish was kept the same (20-30 cm) for both the size classes of IMC. The specific details on body weight, total length and number of individuals stocked, are given in Table 2. All the fishes were procured from the local commercial fish farms. There was no observed mortality for any of the fish species during the experimental duration. Mesocosm with small-fish IMC were sampled at 0-, 30-, 90-, and 120-days time points to measure different biotic and abiotic factors, whereas large-fish ponds were sampled at 0-, 60- and 90-days time points only. The slight irregularity in the sampling intervals was due to restrictions laid by the Government of India over movement during the COVID pandemic. All the sampling protocols were kept the same for both the control and the test ponds, and for both the size classes of IMC.

**Table 2.** Body length (Mean± SE) and weight (Mean± SE) of IMC and catfish during the time of stocking in the mesocosm ponds.

| Fish type |  | Size class | Body length<br>(cm)<br>(Mean± SE) | Size range<br>(cm) | Body weight<br>(g)<br>(Mean± SE) |
| --- | --- | --- | --- | --- | --- |
| IMC | Catla (N=60) | Small-fish | 12.04±0.20 | 10.2-13.8 | 18.1±1.02 |
| (N=300) | Rohu | (10-20 cm) | 14.85±0.40 | 12-17.8 | 30.93±2.34 |
|  | (N=180) |  |  |  |  |
|  | Mrigal(N=60) |  | 10.13±0.28 | 8-13.8 | 13.36±1.02 |
| IMC | Catla (N=60) | Large-fish | 22.67±0.29 | 20-25.2 | 149.35±6.77 |
| (N=300) | Rohu<br>(N=180) | (20-30 cm) | 22.02±0.40 | 20-25.1 | 121.35±7.69 |
|  | Mrigal(N=60) |  | 25.18±0.38 | 21-28 | 140.45±4.89 |
| Catfish<br>(N=80) | Catfish(N=80) | For both<br>small- and<br>large-fish<br>of IMC | 34.21±1.02 | 27.3-41 | 275.4±21.56 |

**Table 3.**
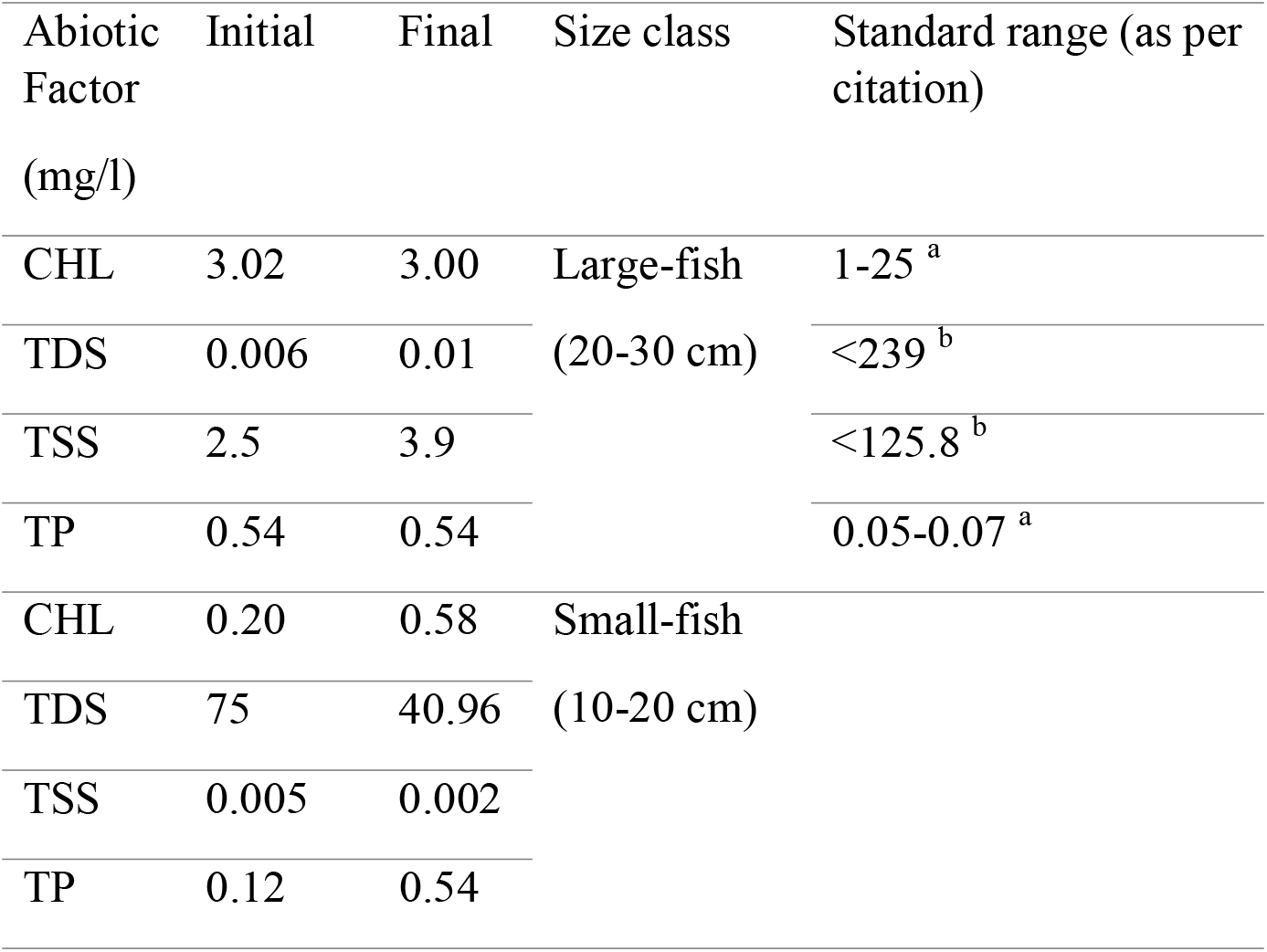
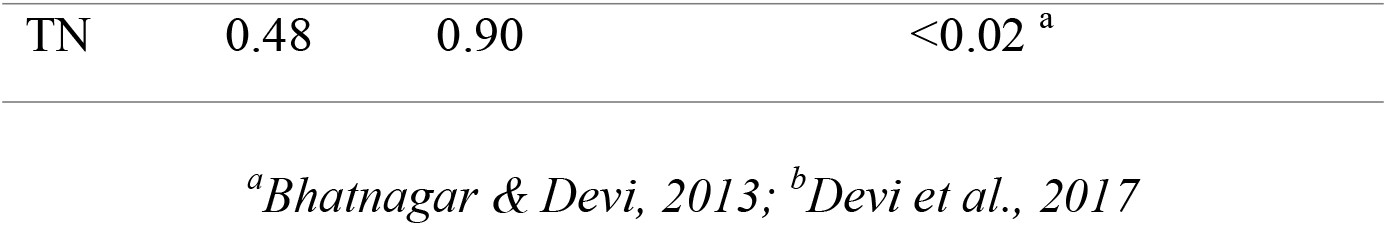
Initial (0 days) and final values of different abiotic factors measured in the test ponds for both large (final sampling was at 90 days) and small-fish (final sampling was at 120 days) of IMC. Standard range indicates values reported in literature as permissible limits for aquaculture ponds within the Indian context. Footnotes indicate the citations used for referring to the permissible limits.

### Sampling of biotic factors

Total length (snout to tail tip) and body weight was used as a proxy to measure growth of an individual fish (Hoover et al., 2013) At each sampling point, fish were sampled for length and weight measurements using drag nets (dimensions, 25*10 m with a mesh size of 12mm) that covered the entire surface area of the ponds. The body length of fish (N=10 individuals of each species) was measured using a measuring scale (1 ft) and weighed on an industrial scale (10 kg scale, Virgo, New Delhi, India). Zooplankton sampling was done during morning hours (7:00-9:00 AM) by collecting 100 ml of water from subsurface (till 50 m below the water surface) (Aminot and Rey, 2000) and filtering into a sterile plastic bottle (100 ml, Tarsons, Kolkata, India) through a plankton net (50-µm mesh, 30 cm diameter). The zooplankton analysis was carried out at Zoological Survey of India (ZSI), Kolkata, India using a stereomicroscope (LEICA M165C, Leica Microsystems, Wetzlar, Germany) and all the samples were transported from the field site to ZSI by adding 1 ml formalin (HiMedia, Mumbai, India) (Besemer, 2009).

A standard size (140*16, diameter*height, mm) sterile plastic petri dish (Tarsons, New Delhi, India) was divided into four equal quadrants, and abundance of zooplanktons at each quadrant was counted using a steromicroscope (10X magnification) for every 50 ml of sample volume (Herrera-Martínez et al., 2017). Zooplankton was identified up to class or subclass (commonly referred to as groups) levels (Doan Dang et al., 2015), and abundance values (Spellenberg and Fedor, 2003) were calculated.

### Sampling of abiotic factors

Abiotic factors, for example, total phosphate (TP), total suspended solids (TSS), total dissolved solids (TDS) including chlorophyll ‘a’(CHL), were measured at each of the sampling points for both the test and the control mesocosms for both the size classes. Total nitrite (TN) for water and carbon and nitrogen ratios (C/N) for soil samples, were measured (all sampling points for TN; 0-, 90-, and 120-days for C/N) for the small-size mesocosms (for both the test and the control ponds) only. All the abiotic factors were measured in 3 technical replicates. The estimation of CHL was carried out by filtering 100 ml of water samples through WHATMAN GF/F glass microfiber filters (pore size 1.2 μm, Millipore, Illinois, USA) (Aminot and Rey, 2000) using a vacuum pump (Millipore, USA), which had 1 ml of acetone (Himedia, Mumbai, India) distributed uniformly prior to filtration in low light conditions. The filter paper was then transferred to a sterile mortar (Pal Surgical and Medical, Ghaziabad, India) and crushed using a pestle to generate a slurry aqueous filtrate. The filtrate was stored in a 2 ml centrifuge tube (Tarsons, New Delhi, India), and covered with aluminum foil and stored overnight in a refrigerator (4°C, Samsung, Seoul, South Korea). Next day, the filtrate was analyzed spectrophotometrically (UV-VIS spectrophotometer, Hinotek, Ningbo, China) at 665 and 750 nm wavelength before [represented by suffix ‘o’ in (1)] and after [represented by suffix ‘a’ in (1)] acidification with 0.2 ml 1% HCL (Himedia, Mumbai, India). The concentration of the CHL (mg/m^3^) was calculated using the equation by Lorenzen (1967):

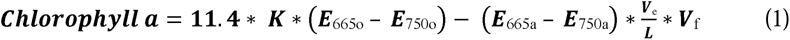

where L = Cuvette light-path (cm), V_e_ = extraction volume (ml), V_f_ = filtered volume (L), E665_o_ and E665_a_ = Absorbance of filtrate at 665 nm before and after addition of HCL, E750_o_ and E750_a_= Absorbance of filtrate at 750 nm before and after addition of HCL, K= R (R-1) = 2.43, here R stands for maximum absorbance ratio of E665_o_ /E 665_a_.

For TSS estimation, 100 ml of pond water was collected and filtered through WHATMAN GF/F glass microfiber filters (pore size 1.2 μm, Millipore, Illinois, USA) using a vacuum pump (Millipore, USA). The filter paper was oven dried (Generic, Ghaziabad, India) and weighed on an analytical balance (Contech, Navi Mumbai, India). The concentration of TSS was determined by the difference in weight of the filter paper, filtered with pond water and that with distilled water (negative control, same volume as sample) divided by volume of the sample (Greenberg et al., 2014). TDS was estimated by evaporation method, where the water samples (100 ml) collected from ponds was filtered through WHATMAN GF/F glass microfiber filters (pore size 1.2 μm, Millipore, Illinois, USA) using a vacuum pump (Millipore, USA), and the fitrate collected in a sterile beaker was subjected to oven (Generic, Ghaziabad, India) drying for an hour. Similarly, filtrate collected after filtering distilled water (negative control, same volume as sample) was also subjected to oven drying for an hour. The dried beakers were weighed (A = pond sample and B = negative control) on an analytical balance (Contech, Navi Mumbai, India), and difference in the weights when divided by the total volume of water sample (V) was referred to as the TDS [TDS= (A-B)/V] (Corwin and Yemoto, 2020).

For analysis of TP, 100 ml of water sample was collected in a sterile reagent bottle (100 ml, Tarsons, New Delhi, India). A standard stock solution was made by mixing 0.1361 g of phosphorus dihydrogen phosphate (Himedia, Mumbai, India) in 100 ml of distilled water containing 1 ml of 9N sulphuric acid (Himedia, Mumbai, India), and working standards (ranging from 0.05 mg/ml-0.005 mg/ml) were prepared from the stock solution (Kaisary et al., 2012). The estimation of total phosphorous was carried out using stannous chloride (Himedia, Mumbai, India), ammonium molybdate (Himedia, Mumbai, India), and sulphuric acid (Himedia, Mumbai, India) solutions, and absorbance was measured at 650 nm using a spectrophotometer (UV-VIS spectrophotometer, Hinotek, Ningbo, China). The amount of phosphorus content in water samples was estimated from the standard curve by measuring the absorbance at the same wavelength. All the reagents and the protocol to estimate TP was followed from Gales et al. (1966) Similarly, for quantification of TN, a stock solution of anhydrous sodium nitrite (Loba Chemie, Mumbai, India) was prepared by dissolving 200 mg of the compound in 100 ml of distilled water. The working standards (ranging from 0.2 mg/ml - 0.002 mg/ml) were prepared from the stock solution of anhydrous sodium nitrite. Further, 1% sulphanilamide (Loba Chemie, Mumbai, India) and 0.1% N-(1-naphthyl)– ethylenediamine dihydrochloride (NEDA) (Loba Chemie, Mumbai, India) were prepared and stored in amber glass bottles. Both the reagents were added to blank, standards and water samples (100 ml). A standard calibration curve was prepared using a range of concentrations of working standards along with their corresponding absorbance values at 540 nm and was used to quantify the unknown nitrite concentration of the pond water samples (Kaisary et al., 2012).

Soil samples (approx. 100g) were collected using a corer (PVC, 42.10 mm diameter, 30 cm length). The collected soil samples were sun-dried at the field site and then transferred to the laboratory in 100 ml plastic containers (Pride Homeware, Kolkata, India). The soil samples were homogenized and dried for 24 hours in a hot air oven (Vindish Instruments Private Limited, Ahmedabad, India) at 55°C followed by physical removal of twigs, or stones using a pair of forceps. The dried soil samples were crushed into a fine powder, using mortar and pestle (Pal Surgical and Medical, Ghaziabad, India). The powdered samples were weighed and 2-3 g of samples were placed into a 15 ml centrifuge tube (Tarsons, New Delhi, India), which was subjected to decarbonation in order to remove the inorganic carbon. Briefly, during the decarbonation process, powdered samples were mixed twice with 10 ml of 1N HCL (Merck, New Jersey, USA) by vortexing (Spinx vortex shaker, Tarsons, New Delhi, India) for 30 seconds and the top aqueous layer was dis-carded. Then the residue was placed on a hot air oven for an hour to remove acid from the soil samples. Following this, the sample was rinsed with 10 ml of Milli-Q water (Merck, New Jersey, USA), and centrifuged at 2500 rpm for 5 minutes using a table-top centrifuge (Remi Elektrotechnik, Maharashtra, India). The Milli-Q rinse was repeated until the solu-tion turned neutral, which was confirmed by a litmus test. Post decarbonation, the soil samples were placed overnight in a hot air oven at 55°C. The C/N ratio was analyzed in elemental analyzer (Elementar, Model: Vario Isotope select, Germany, Europe) using combustion method (Banerji et al., 2021). The amount of carbon and nitrogen content in the soil samples were estimated from the calibrated standard curve prepared from High Organic Sediment Sample (HOSS) with a known carbon (6.72%) and nitrogen (0.5%) content (Banerji et al., 2021). A standard curve was prepared using different weights of HOSS starting from 1.16 -100.60 mg. Samples were weighed upto 100 mg and placed in a tin cup for analysis. The quantity of carbon to nitrogen ratio in the unknown samples was measured from the standard curve (Banerji et al., 2021). Since the carbon to nitrogen values are represented as ratios, we did not perform any statistical analysis and the C/N values are given in Supplementary Table 1. All samples were analyzed in replicates for each time point (0-, 90-, 120-days).

### Statistical analysis

Small- and large-size mesocosm experiments were performed during different times of the year (calendar year corresponding to different seasons or differences in ambient temperature), thus, we analyzed the data separately instead of including them in a single statistical model. All the data was first checked for normality assumption, and if the data followed normality (raw data or after log or log x+1 transformations), parametric analysis using linear mixed effect (LME, random effects were included to avoid pseudoreplication as the same ponds were sampled over time) model with maximum likelihood method was used. The treatments (with two levels, control and test) and days (sampling points starting from 0-days to 90- or 120-days) were included as independent variables, and fish morphometrics (length and weight), zooplankton abundance and hydrological parameters were included as a dependent variables. If normality assumption was not met, we used Kruskal-Wallis test for comparisons over time (sampling points) for a given mesocosm (separately for test and control ponds), and Mann-Whitney-Wilcoxon test to compare between overall values (pooled across sampling points) for test and control ponds. The abiotic factors, TDS and TP, in large-size IMC ponds had a significant difference (P<0.05, Mann-Whitney-Wilcoxon test) between control and test mesocosms at the 0-days or first sampling point. Thus, for TDS and TP only, relative values were taken into account for statistical analyses, for example, [TP value at 60-days sampling point–TP value at 0-day sampling point, and likewise for all sampling points]. For all statistical tests, appropriate post-hoc analysis was carried out only if there was a significant difference in the overall values (pooled across sampling points) between the control and the test ponds. However, if there was a significant difference with respect to time (sampling points), post-hoc pairwise analyses between sampling points (0 versus 90 or 120 days) were not included as that is not the primary focus of the manuscript. Tables 4 and 5 lists the type of statistical tests for each parameter. All the statistical analysis was done on the R program, ver.3.2.1. Significance level was fixed at P< 0.05 for all the analyses.

**Table 4.**
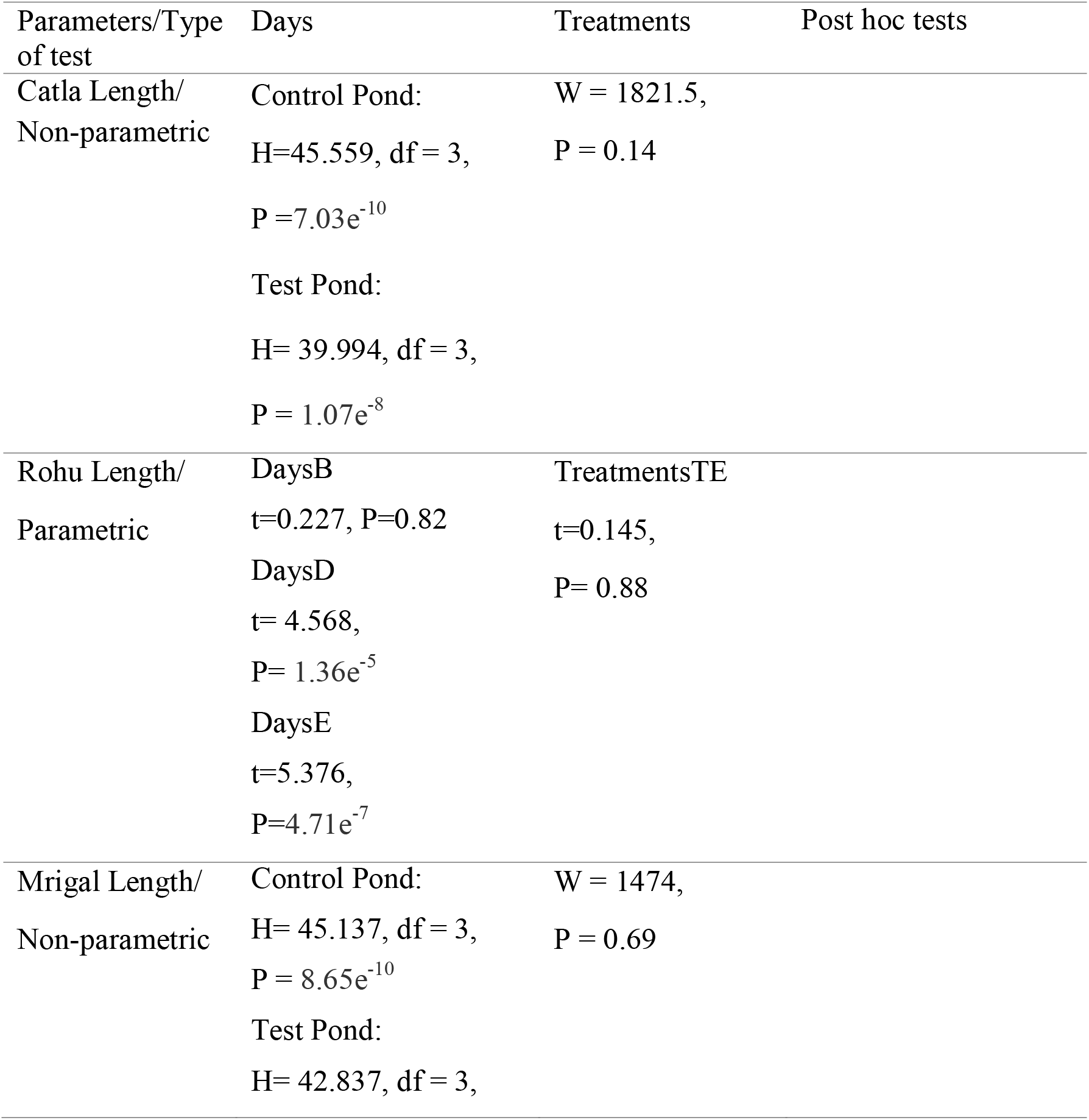
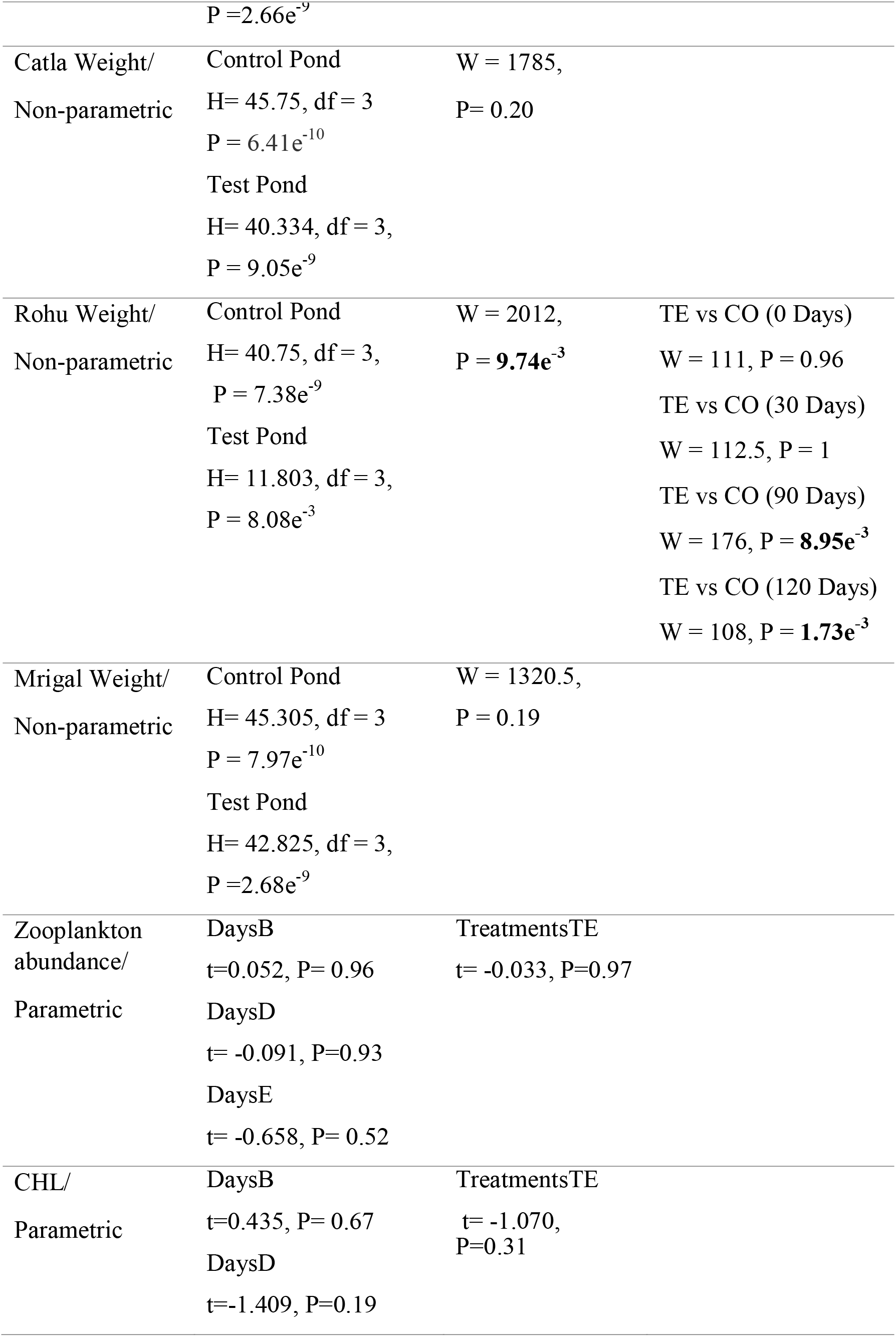
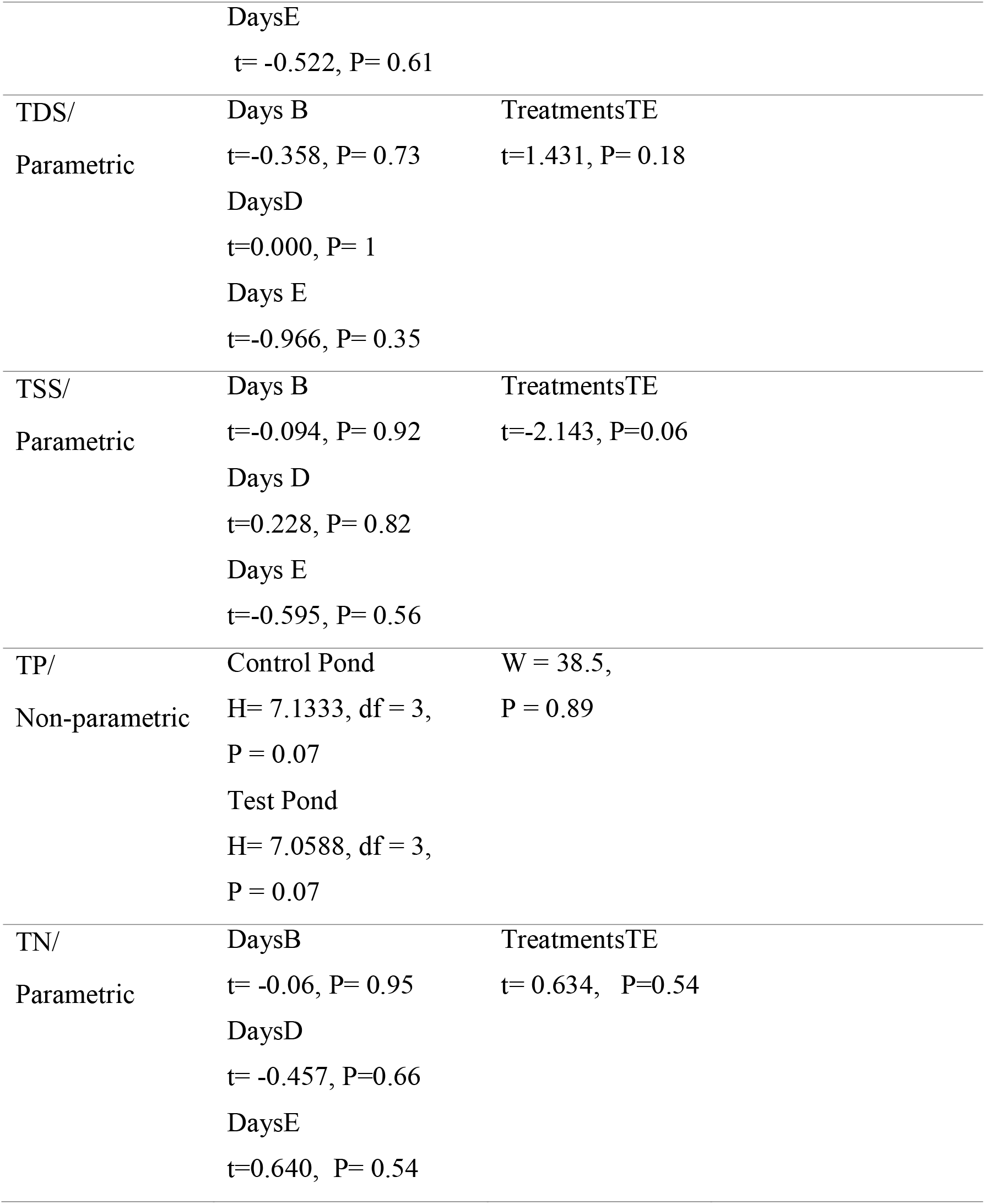
Statistical results of the mesocosm experiments for the small-fish of IMC. Results for treatment*days are not shown as none of the factors in parametric analysis (treatment or days) gave significant results. All significant (P<0.05) values while comparing test to control ponds are indicated in bold. Post hoc tests are only shown for significant results in the treatment column while comparing control versus test ponds.

| Parameters/Type of test | Days | Treatments | Post hoc tests |
| --- | --- | --- | --- |
| Catla Length/<br>Non-parametric | Control Pond:<br>H=45.559, df = 3,<br>P = 7.03e<sup>-10</sup><br>Test Pond:<br>H= 39.994, df = 3,<br>P = 1.07e<sup>-8</sup> | W = 1821.5,<br>P = 0.14 |  |
| Rohu Length/<br>Parametric | DaysB<br>t=0.227, P=0.82<br>DaysD<br>t= 4.568,<br>P= 1.36e<sup>-5</sup><br>DaysE<br>t=5.376,<br>P=4.71e<sup>-7</sup> | TreatmentsTE<br>t=0.145,<br>P= 0.88 |  |
| Mrigal Length/<br>Non-parametric | Control Pond:<br>H= 45.137, df = 3,<br>P = 8.65e<sup>-10</sup><br>Test Pond:<br>H= 42.837, df = 3, | W = 1474,<br>P = 0.69 |  |
|  | P = 2.66e<sup>-9</sup> |  |  |
| Catla Weight/<br>Non-parametric | Control Pond<br>H= 45.75, df = 3<br>P = 6.41e<sup>-10</sup><br>Test Pond<br>H= 40.334, df = 3,<br>P = 9.05e<sup>-9</sup> | W = 1785,<br>P= 0.20 |  |
| Rohu Weight/<br>Non-parametric | Control Pond<br>H= 40.75, df = 3,<br>P = 7.38e<sup>-9</sup><br>Test Pond<br>H= 11.803, df = 3,<br>P = 8.08e<sup>-3</sup> | W = 2012,<br>P = **9.74e<sup>-3</sup>** | TE vs CO (0 Days)<br>W = 111, P = 0.96<br>TE vs CO (30 Days)<br>W = 112.5, P = 1<br>TE vs CO (90 Days)<br>W = 176, P = **8.95e<sup>-3</sup>**<br>TE vs CO (120 Days)<br>W = 108, P = **1.73e<sup>-3</sup>** |
| Mrigal Weight/<br>Non-parametric | Control Pond<br>H= 45.305, df = 3<br>P = 7.97e<sup>-10</sup><br>Test Pond<br>H= 42.825, df = 3,<br>P = 2.68e<sup>-9</sup> | W = 1320.5,<br>P = 0.19 |  |
| Zooplankton<br>abundance/<br>Parametric | DaysB<br>t=0.052, P= 0.96<br>DaysD<br>t= -0.091, P=0.93<br>DaysE<br>t= -0.658, P= 0.52 | TreatmentsTE<br>t= -0.033, P=0.97 |  |
| CHL/<br>Parametric | DaysB<br>t=0.435, P= 0.67<br>DaysD<br>t=-1.409, P=0.19 | TreatmentsTE<br>t= -1.070,<br>P=0.31 |  |

|  |  |  |
| --- | --- | --- |
|  | DaysE |  |
|  | t= -0.522, P= 0.61 |  |
| TDS/<br>Parametric | Days B<br>t=-0.358, P= 0.73<br>DaysD<br>t=0.000, P= 1<br>Days E<br>t=-0.966, P= 0.35 | TreatmentsTE<br>t=1.431, P= 0.18 |
| TSS/<br>Parametric | Days B<br>t=-0.094, P= 0.92<br>Days D<br>t=0.228, P= 0.82<br>Days E<br>t=-0.595, P= 0.56 | TreatmentsTE<br>t=-2.143, P=0.06 |
| TP/<br>Non-parametric | Control Pond<br>H= 7.1333, df = 3,<br>P = 0.07<br>Test Pond<br>H= 7.0588, df = 3,<br>P = 0.07 | W = 38.5,<br>P = 0.89 |
| TN/<br>Parametric | DaysB<br>t= -0.06, P= 0.95<br>DaysD<br>t= -0.457, P=0.66<br>DaysE<br>t=0.640, P= 0.54 | TreatmentsTE<br>t= 0.634, P=0.54 |

**Table 5.**
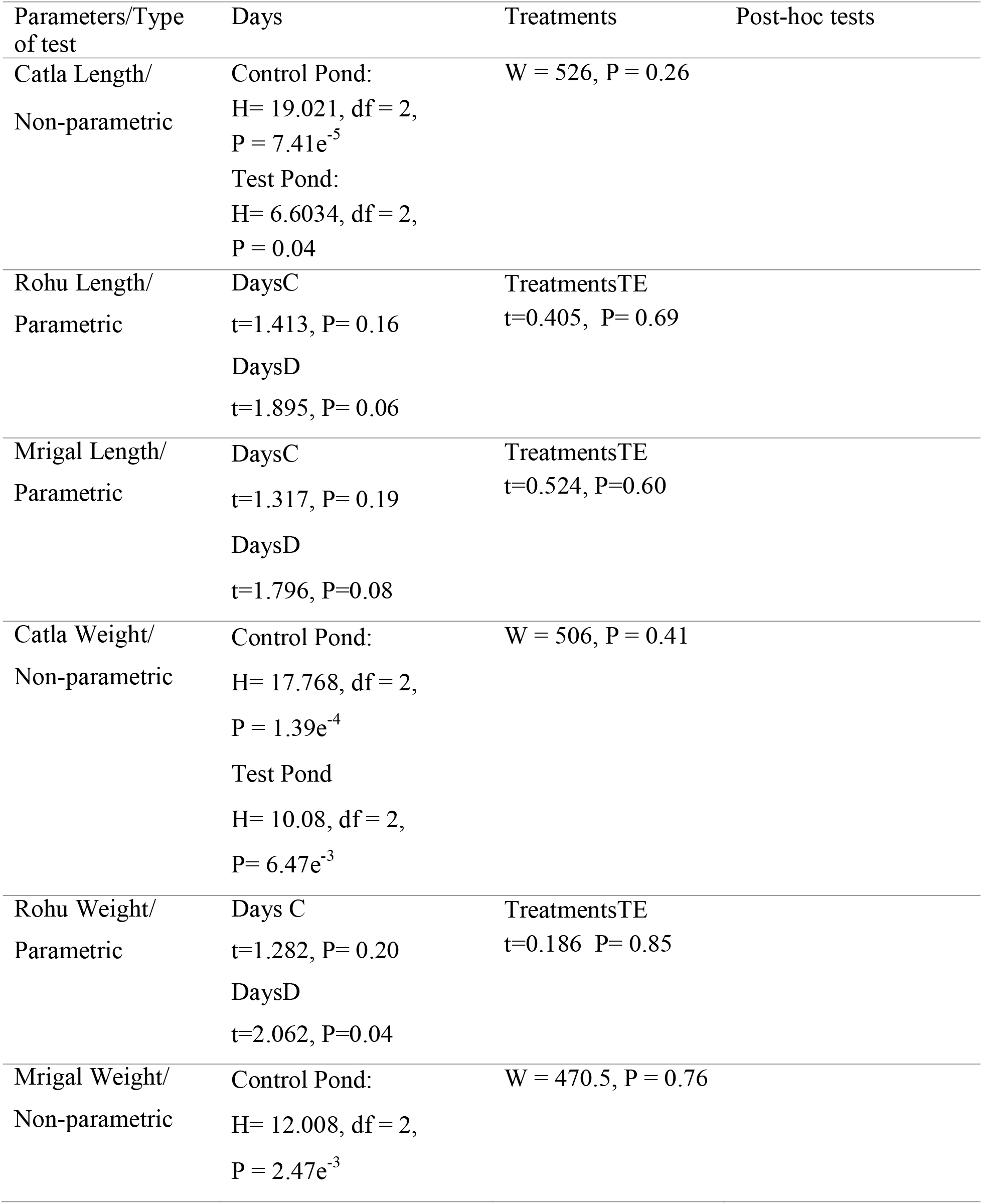
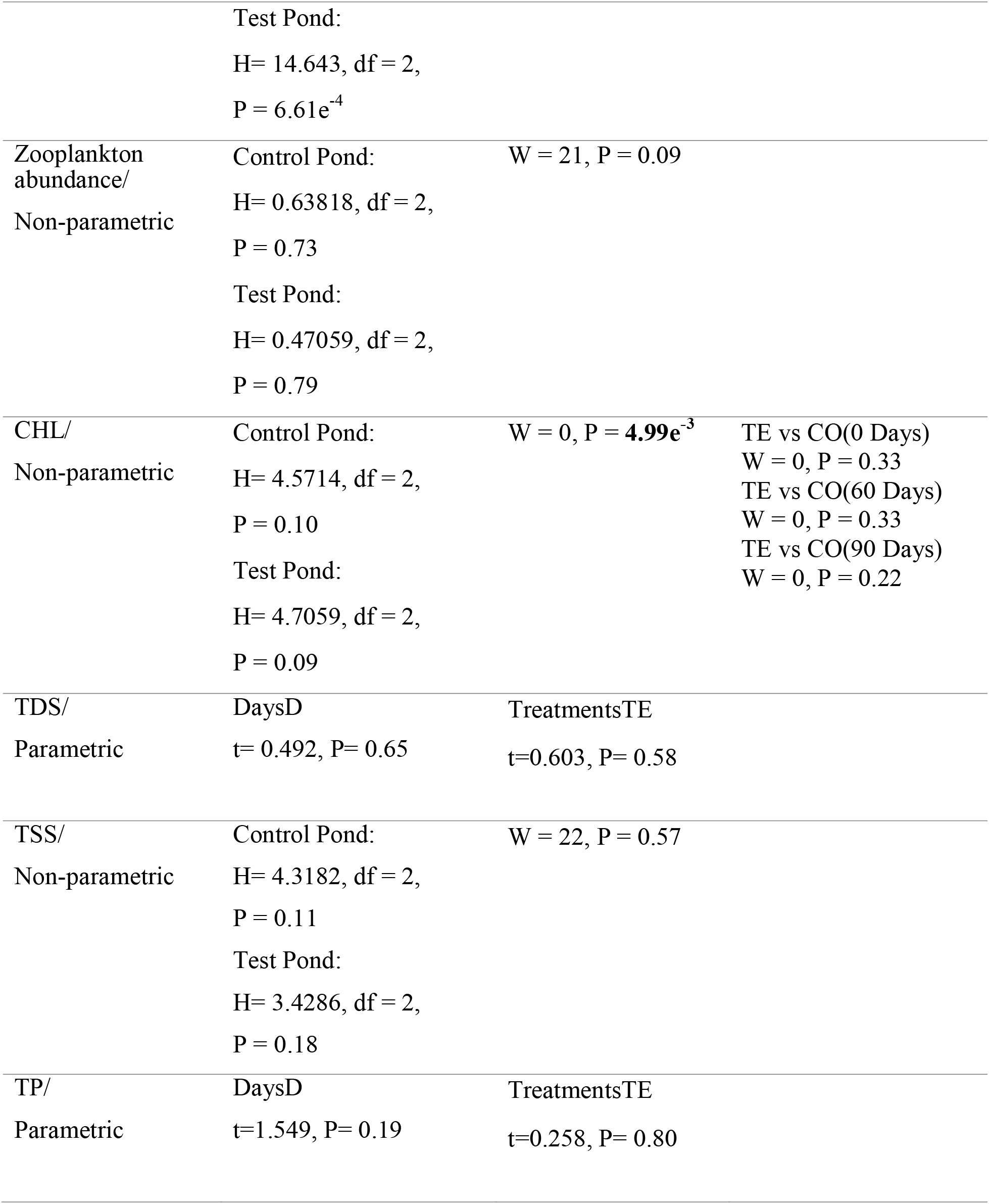
Statistical results of the mesocosm experiments for the large-fish of IMC. Results for treatment*days are not shown as none of the factors in parametric analysis (treatment or days) gave significant results. All significant (P<0.05) values while comparing test to control ponds are indicated in bold. Post hoc tests are only shown for significant results in the treatment column while comparing control versus test ponds.

## RESULTS

All hydrological factors, for example, CHL, TSS and TDS, except TP and TN for the small-fish of IMC, and TP for the large fish of IMC, were within the standard permissible range as reported in literature for aquacultural ponds and freshwater habitats within India (Table 3). The weight of rohu declined significantly (W=2012, P=9.74e^-3^, Mann-Whitney-Wilcoxon test) (Fig. 1, Table 4) in the test pond when compared to control, and the reduction in weight was significant at 90-(P=8.95e^-3^, post-hoc Mann-Whitney-Wilcoxon test) and 120-days (P=1.73e^-3^, post-hoc Mann-Whitney-Wilcoxon test) sampling points only. Apart from reduced weight in small-size rohu, there were no other significant differences (P>0.05) in the measured biotic factors between control and test ponds for both the small-(Table 4) and the large-size classes (Table 5) of IMC. The lengths (Mean±SE) of large-size rohu, catla and mrigal at final sampling point were 28.78 ± 0.85 cm in control and 27.01 ± 0.47 cm in test, 26.69 ± 1.11 in control and 27.19 ± 1.23 cm in test, and 30.75 ± 1.10 cm in control and 29.33 ± 0.95 cm in test ponds, respectively. For the small-size, the final lengths were 21.78 ± 0.42 cm in control and 16.88 ± 0.97 cm in test for rohu, 24.78 ± 0.53 in control and 21.36 ± 0.91 cm in test ponds for catla, and 25.17 ± 1.08 in control and 22.99 ± 1.12 cm in test ponds for mrigal. For the abiotic factors, only CHL showed significant differences between control and test ponds for the large-fish of IMC (W = 0, P = 4.99e^-3^, Mann-Whitney-Wilcoxon test), however, no significant differences (P>0.05) were observed in post-hoc pairwise comparisons (Table 5).

**Fig. 1.**
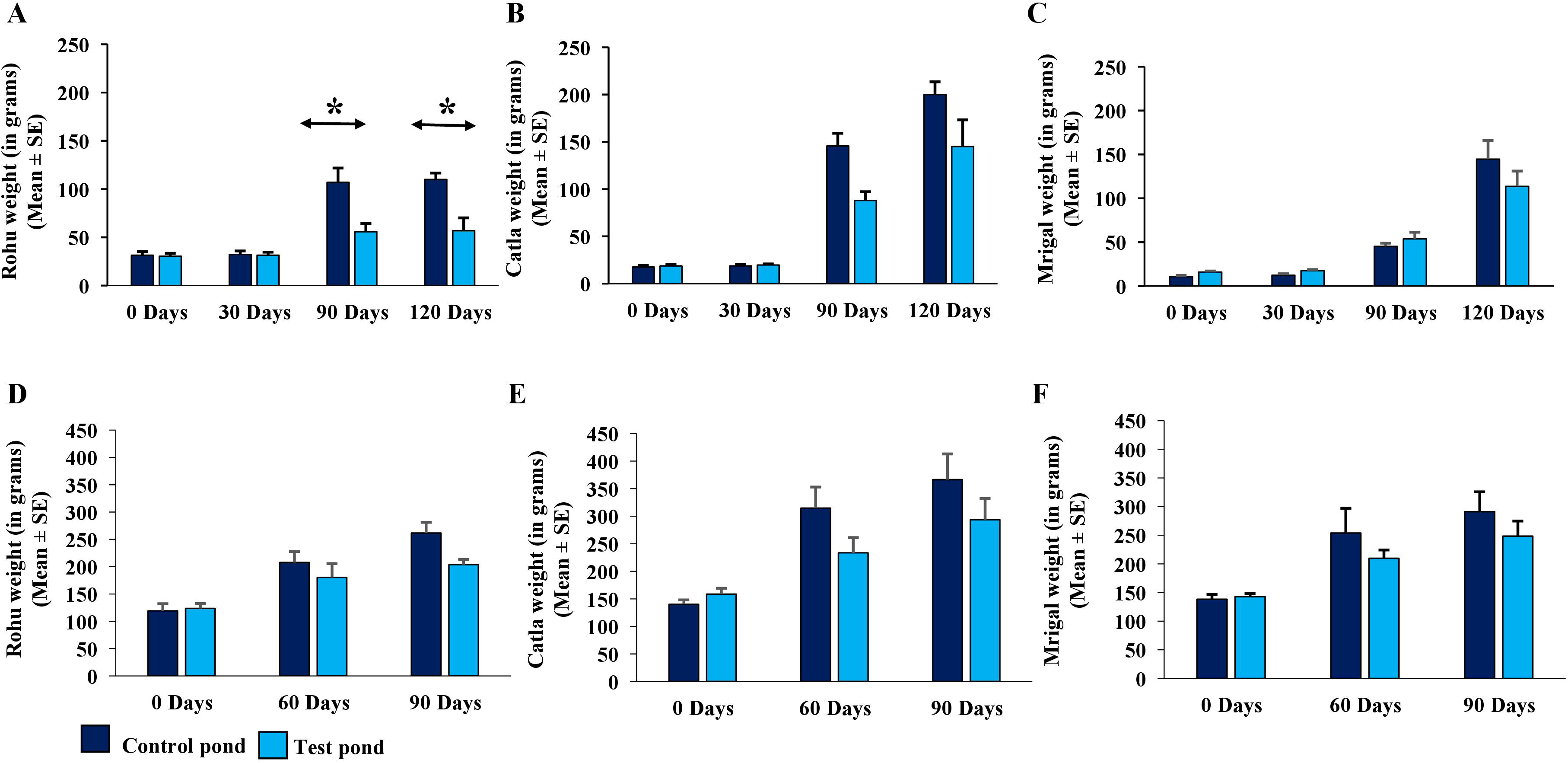
Weight (Mean±SEM) of IMC in control and test ponds. Panels A, B and C indicate weights for small-size (10-20 cm) and panels D, E, and F represent weights for large-size (20-30 cm) of IMC. *indicates significant differences between control and test ponds for a particular sampling point.

With regards to the zooplankton classification and abundance (Mean± SE percentages pooled across sampling points, n=4), copepods (90.91±4.24) and cladocera (9.09 ±4.24,) were the predominant families for the small-fish mesocosms. However, for the large-fish (Mean± SE percentages pooled across sampling points, n=3), copepods (29.61±8.34), cladocera (17.35±5.02) and rotifers (53.03±13.35) were the predominant groups. With regards to soil analysis for small-size class only, the control pond had a mean (n=6, 2 assay replicates for each time point) C/N ratio of 8.38 whereas the test pond had a mean (n=6, 2 assay replicates for each time point) C/N ratio of 9.29. The C/N values for each time point for both the control and the test ponds are given in Supplementary Table 1.

## DISCUSSION

Our results demonstrate that the impact of an exotic fish species can vary depending on the developmental stage (as indicated by body length in our study) of the native species. Further, certain native species, for example, rohu in our study, appeared to be more sensitive towards the presence of an exotic when compared to other IMC species. We also found that the sailfin catfish had no significant impact on the abiotic factors, and on the abundance of zooplankton in the test mesocosms for both the size classes of the native species. Thus, it can be speculated that reduced growth of rohu in small-size mesocosm could be due to behavioural competition between the native and the exotic species during the time of foraging.

The present study shows that sailfin catfish significantly impacted the growth of rohu in small-size mesocosm. This is in agreement with several studies, where smaller size classes or juveniles have higher vulnerability towards exotic invasions when compared to adult ones (Hrabik et al., 1998; Mills et al., 2004; Ayala et al., 2007). For example, Hrabik et al. (1998) demonstrated the impact of the introduced rainbow smelt *Osmerus mordax* over native cisco fish species *Coregonus artedii* in an oligotrophic lake in Wisconsin, USA. The authors conducted a long-term survey (1981-1994), and showed a significant decline in abundance of cisco population, post-introduction of rainbow smelt. The reduction in native cisco fishes was due to excessive predation by smelt, particularly over the young of the year and the juveniles. Similarly, Mills et al. (2004) demonstrated the impact of an introduced species, western mosquitofish *Gambusia affinis*, over two size classes [small (9-13 mm) and large (23-28 mm)] of juvenile least chub *Lotichthys phlegethontis*, a native species in Utah, USA. The study showed that presence of mosquitofish significantly impaired growth and survival of the small-size least chub when compared to the large-size ones. The authors speculated that the impact on smaller individuals could be due to competition for food in presence of the exotic species and/or a higher rate of predation by the exotic mosquitofish. However, the impact exerted by an exotic species may not always be specific towards certain developmental stages of the natives, and may equally impact different stages of an organism, as well. McIntosh, (2000) studied the impact of the introduced brown trout *Oncorhyncus mykiss* over the native population of common galaxias *Galaxias vulgaris* belonging to different size classes in freshwater lakes of New Zealand. The study showed that large body (>150 mm) trout significantly predated common galaxias irrespective of their development stages (small, juvenile and adult attaining a length of 90-150 mm). Whether an exotic is impactful all across or only towards certain developmental stages needs systematic assessments because prey-predator interactions and preferences for different food items vary largely along the life stages of an organism (Julius et al., 2012; Sampson et al., 2009).

Our mesocosm experiments showed that only one of the native species of IMC, rohu, responded significantly in terms of reduced growth towards the presence of the Amazon sailfin catfish. Similar results have been documented by several studies highlighting that certain species are more sensitive or vulnerable towards the presence of exotics when compared to others within an experimental setup (Wahab et al., 1995; Pendleton et al., 2014) or under natural conditions (McIntosh, 2000; Moi et al., 2021; Crowl et al., 1992). For example, Pendleton et al., (2014) conducted mesocosm experiments to demonstrate the impact of an exotic predatory fish, piranha *Serrasalmus marginatus* over a native fish species, pike characid *Acestrorhynchus lacustris* in tropical floodplains of Upper Parana River, Brazil. The study showed that out of the 18 species that were available in the floodplains, only the pike populations declined in presence of the piranha, which in turn impacted the phosphorus cycle, as the pikes are known to sequester phosphorus from the soil into the freshwaters (Small et al., 2011). Our study also demonstrated that sailfin catfish had no significant impact over the growth of the two of the IMC species, catla and mrigal. Similar results have been documented by Chaichana and Jongphadungkiet (2012), which demonstrated that Amazon sailfin catfish had no significant impact on the growth of the bighead catfish *Clarius macrocephalus*, native to Thailand. Whether an introduced exotic is negatively impacting a single (Wahab et al., 1995; Pendleton et al., 2014; Mazzeo et al., 2010) or multiple species (Maezono et al., 2005; Carey & Wahl, 2010) or is not impactful at all (Chaichana and Jongphadungkiet, 2012), needs experimental monitoring at the community level for a particular ecosystem that faces a biological invasion.

The impact of an introduced species heavily depends on the resilience of an ecosystem (Thoms et al., 2018), thus, mesocosm experiments need to be conducted over time to understand whether a given ecosystem can tolerate the impact of a stressor (introduced species) (Aminot and Rey, 2000; Callaway et al., 2005) or not (Scheffer, 1989). Long-term studies (Phelps et al., 2017; Chick et al., 2020) are regarded as valuable when compared to short-term or one-time assessments (Schrank et al., 2003; Sampson et al., 2009) in terms of understanding dynamics of a particular ecosystem over time (pre- and post-introductions of exotics, and/or over time since introduction of an exotic) (Strayer et al., 2006). For example, Beisner et al., (2003) demonstrated the impact of introducing rainbow smelt over the native fish and zooplankton communities in two lakes, Crystal and Sparkling, in Wisconsin, USA. The authors compared the impact of smelt between short-term (2 weeks) and long-term (1981-1999 for Crystal lake, 1981-1983 for Sparkling lake) assessments. The short-term study on both the lakes demonstrated exotic smelt had no impact on zooplankton community as they rarely predated upon zooplankton when compared to the native planktivorous fishes, cisco and yellow perch *Perca flavescens*. However, evaluating the ecological communities over a longer time showed the exotic smelt was indirectly impacting the zooplanktons in both the lakes by directly predating upon the native planktivore fishes. Time series analysis showed that predation by exotic smelt over the young of the year of the native fishes (cisco and perch) resulted in alteration of the zooplankton community structure for both the lakes. Apart from the biotic factors, exotic species can also indirectly impact the native species by altering the abiotic factors (hydrological and soil parameters) of an ecosystem (Drenner et al., 1997).

The abiotic factors measured during our mesocosm study were within the range of values that have been documented for most aquaculture ponds and freshwater habitats within the Indian context (Bhatnagar & Devi, 2013; Devi et al., 2017). Thus, the established mesocosms closely mimicked the Indian open waters, which has been invaded by the catfish. However, values for TN (both test and control for small-size) and TP (both test and control ponds for large- and small-size) were the exceptions. The higher range of TN could be due to precipitation (Lin et al., 2002) whereas for TP, rainwater runoff (Kiran, 2010) and dead organic matter from leaves (Chamberlain et al., 2001) could be the sources that increased phosphorus in the mesocosms. Interestingly, most studies have shown that exotic species impact the native ecosystems by habitat modification thereby altering the hydrological and soil parameters that impair the survival and growth of other native species within an ecosystem (Roberts et al., 1995; Tan et al., 2010). However, this is in contrast to our study that showed no significant impact of the sailfin catfish over the abiotic factors of the test mesocosms. Similar results have also been documented for several exotic species (Venturelli & Tonn, 2005; Tsang & Dudgeon, 2021). For example, Tsang & Dudgeon, (2021) assessed the impact of two exotic fish species, mosquitofish and guppy, over population of native fish species, rice fish *Oryzias curvinotus* and half banded barb *Puntius semifasciolatus*, invertebrate abundance, and biomass of phytoplankton and periphyton, and several abiotic factors (nitrite, nitrate and ammonia concentration) using a mesocosm approach in Hong Kong, China. The study demonstrated that the exotic fish contributed towards decline in invertebrate abundance, which in turn impacted the population of native fishes. However, there were no significant differences in the abiotic factors between control (absence of exotic) and test (presence of exotic) mesocosms. The authors speculated that high predation rate by the exotic species on invertebrate larvae could have led the native fish species towards food deprivation and eventually declining their population sizes (Tsang & Dudgeon, 2021). Thus, this prioritizes the need for behavioral studies, for example, measuring aggression (an indicator of competition) in exotics in presence of the native species under different contexts, feeding, foraging and nest building. Apart from ecological interests, the introduction of an exotic species raises further concerns when the impacted native species belong to economically or commercially valuable food fish category.

IMC that we used for our study are economically valuable food fish (Dwivedi et al., 2004) and contributes to about 58% of aquaculture production in India with a gross commercial value of ∼$3.11/kg (as reported in 2021) (Karnatak et al., 2021) Among the IMC species, rohu is the most preferred aquaculture species for its considerably higher market price in comparison to catla and mrigal (Asaduzzaman et al., 2010; Rasal & Sundaray, 2020). Rohu is primarily a column feeder mostly predating upon planktons and periphytons (Majumder et al., 2020). Wijethunga and Epa, (2008) used gut content analysis to report 66% of dietary overlap between exotic sailfin catfish and indigenous species within freshwaters of Sri Lanka that predominantly included species like catla and rohu. Similar results have been documented by other studies (Mendoza-Carranza et al., 2010, Hubilla et al., 2008), as well. Moreover, a mesocosm-based study by Seshagiri et al., (2021) demonstrated that presence of sailfin catfish diminished the IMC production by 22.92%. Further, the study speculated that exotic sailfin catfish contributed to food chain disruption due its voracious feeding habits over benthic algae, and periphyton, thus, possibly giving huge competition to the native IMC species in terms of food availability and foraging success. This is in support of our results that demonstrated no significant impact of the exotic catfish on zooplankton abundance and abiotic factors of the test mesocosms, and thus, speculating that the impact of catfish on growth of small rohu could be due to feeding competition between the two species. Though, our study did not measure algae and periphyton, and thus, needs to be included in future work. Given the anecdotal evidence on behavioural competition between exotic catfish and native rohu species, it will be interesting to conduct observations on the feeding behaviours of the two species in presence of each other, and thus, will provide insights into the mechanisms by which the exotic is impacting the natives.

## CONCLUSIONS

Overall, the study showed that the Amazon sailfin catfish negatively impacted the growth of small-size rohu, post 90-days of stocking. Since there were no significant changes in abiotic factors between control and test ponds, the weight loss in small rohu can be attributed to behavioral competitions between native and exotics for food resources. Hence, future work should investigate foraging behavior and competition between the native rohu and the exotic catfish to better understand the negative impact. A multi-pronged approach, including several native species and measurements of abiotic factors, is desired to understand the dynamics of an ecosystem post-invasion of an exotic. Our study included several such readouts for both biotic and abiotic measurements; however, we did not include a few parameters, for example, benthic algal growth, parasitic load and gut microbiome analysis in native species. Such inclusions may further improve the robustness of the impact assessment analysis. Moreover, such ecological evaluations need to be conducted for combinations of different developmental stages (for both native and exotic species), and across different spatial (for different ecosystems having diverse structure and functions, which have been invaded by the same exotic species) and temporal (time lapsed since introduction of the exotic for a given ecosystem) scales to infer robust conclusions. Such kinds of ecological assessments will provide insights towards better management (Hussan et al., 2021) of the aquatic ecosystems, which are severely threatened by introductions of several exotic species including both vertebrates and invertebrates.

## ACKNOWLEDGMENTS

We would like to thank all the staff at both Rahara and Kalyani field stations of the Indian Council of Agricultural Research (ICAR), Central Institute of Freshwater Aquaculture (CIFA), for help and cooperation during the mesocosm experiments. We are thankful to Dr. Chitra Jayapalan from Zoological Survey of India (ZSI), Kolkata for guidance and support in plankton analysis. We would also like to thank Dr. Ravi Bhushan, Physical Research Laboratory (PRL), Ahmedabad for C/N analysis of soil samples. SM acknowledges DST-SERB, Government of India for providing fellowship during the duration of the research project.

## FUNDING

This work has been supported by Science and Engineering Research Board (SERB), Department of Science and Technology (DST), Government of India (AU/DBLS/SERB-CRG/2018/001909/2019-20/02)

## CONFLICT OF INTEREST

The authors declare that they have no conflicts of interest.

## AUTHOR CONTRIBUTIONS

Ratna Ghosal, Suman Mallick, Jitendra Kumar Sundaray and Ajmal Hussan: conceptualization and design of the experiment; Jitendra Kumar Sundaray, Ratna Ghosal, Ajmal Hussan and Suman Mallick: methodology; Suman Mallick and Ajmal Hussan: formal analysis; Suman Mallick and Ratna Ghosal: draft preparation; Ratna Ghosal, Jitendra Kumar Sundaray and Ajmal Hussan: Review and editing; Suman Mallick and Ratna Ghosal: data visualization; Ratna Ghosal, Jitendra Kumar Sundaray and Ajmal Hussan: validation; Ratna Ghosal and Jitendra Kumar Sundaray: funding acquisition. All authors have read and agreed to the published version of the manuscript.

## Notes

### Competing Interest Statement

The authors have declared no competing interest.

