## Supplemental data for "Ecological assessment of the Amazon sailfin catfish (*Pterygoplichthys* species) within the Indian freshwaters: a mesocosm-based approach"

**Supplementary Table 1**. C/N ratio of soil samples corressponding to different sampling points from both control and test mesocosms for small-size class of IMC

| Sample type | Sampling points | C/N ratio (Mean) |
| --- | --- | --- |
| Control Pond | 0-days | 7.595 |
|  | 90-days | 7.905 |
|  | 120-days | 9.65 |
| Test Pond | 0-days | 7.985 |
|  | 90-days | 10.235 |
|  | 120-days | 9.65 |
